# Chromatin fiber breaks into clutches under tension and crowding

**DOI:** 10.1101/2021.11.16.468645

**Authors:** Shuming Liu, Xingcheng Lin, Bin Zhang

**Affiliations:** Department of Chemistry, Massachusetts Institute of Technology, Cambridge, MA, USA

## Abstract

The arrangement of nucleosomes inside chromatin is of extensive interest. While in vitro experiments have revealed the formation of 30 nm fibers, most in vivo studies have failed to confirm their presence in cell nuclei. To reconcile the diverging experimental findings, we characterized chromatin organization using a near atomistic model. The computed force-extension curve matches well with measurements from single-molecule experiments. Notably, we found that a dodeca-nucleosome in the two-helix zigzag conformation breaks into structures with nucleosome clutches and a mix of trimers and tetramers under tension. Such unfolded configurations can also be stabilized through trans interactions with other chromatin chains. Our study supports a hypothesis that disordered, in vivo chromatin configurations arise as folding intermediates from regular fibril structures. We further revealed that chromatin segments with fibril or clutch structures engaged in distinct binding modes and discussed the implications of these inter-chain interactions for a potential sol-gel phase transition.

## Introduction

Eukaryotic genomes are packaged into nucleosomes by wrapping DNA around histone proteins. While the structure of a single nucleosome has been extensively characterized, ^1–3^ the organization for a string of nucleosomes, i.e., chromatin, remains debatable.^4–6^ Regular, fibril configurations are commonly observed in experiments that study chromatin materials extracted from the nucleus. ^7–10^ The invention of in vitro reconstituted nucleosome arrays with strong-positioning DNA sequences ^11^ helped to remove sample heterogeneity in nucleosome spacing and made possible the determination of high-resolution structures.^12–16^ However, despite the large amount of evidence supporting their formation in vitro, fibril structures are rarely detected by in vivo experiments that have managed to characterize chromatin at a fine resolution.^17–20^ Therefore, their biological relevance has been questioned, and chromatin organization inside the nucleus remains controversial.

It is worth noting that the nuclear environment is rather complex. In addition to interactions among nucleosomes, many factors, including tension and crowding, can impact chromatin organization. Chromatin is known to associate with various force-generating protein molecules involved in transcription and nucleosome remodeling.^21–25^ Furthermore, chromatin is often attached to the nuclear envelope and other liquid droplet-like nuclear bodies.^26–29^ Dynamical fluctuations in these nuclear landmarks could exert forces on chromatin as well.^30,31^ Finally, local nucleosome density can be quite high, especially in heterochromatin regions.^32^ Such a crowded environment could lead to cross-chain contacts that might compete with interactions stabilizing single-chain conformations. ^17^ Therefore, both tension and crowding could destabilize the most stable configuration for isolated chromatin.

Unfolding from the fibril configuration may provide a simple yet appealing explanation for the lack of regular chromatin organization inside the nucleus. Recently, we explored the folding landscape and pathways of a tetranucleosome, i.e., a minimal fibril unit.^33^ Schalch et al. captured the tetra-nucleosome in a two-helix zigzag conformation with X-ray crystallography.^34^ Unfolded tetra-nucleosome configurations from the crystal structure indeed resemble those reported by in vivo studies, including the formation of trinucleosome motifs.^35^ Whether the same conclusion can be generalized to longer chromatin segments and whether tension and crowding stabilize unfolding intermediates remain to be shown.

Chromatin unfolding has indeed been studied extensively with various techniques. ^36,37^ Single-molecule force spectroscopy is a powerful tool for characterizing chromatin organization under tension.^38–41^ Force-extension curves at low-force regimes are particularly informative regarding inter-nucleosomal interactions.^42^ Single-molecule Förster resonance energy transfer is another popular technique for probing nucleosome contacts and chromatin conformational dynamics. ^43–46^ Mesoscopic modeling has also been frequently used to interpret experimental data with structural details. ^47–54^ However, because of the experimental techniques’ low resolution and assumptions on nucleosome-nucleosome interactions introduced in computational models, the exact conformations of unfolded chromatin have not reached a consensus and necessitates further investigations.

We perform computer simulations of a 12-nucleosome-long chromatin segment (12mer) to investigate chromatin unfolding under tension and crowding. Near-atomistic representations are adopted for protein and DNA molecules to capture their interactions with physical chemistry potentials at high resolution. Using a combination of enhanced sampling techniques and machine learning, we show that the computed force-extension curve agrees well with results from single-molecule force spectroscopy experiments. ^55^ Our simulations support chromatin unfolding under tension proceeds through intermediate structures with nucleosome clutches, i.e., configurations that have been directly observed via super-resolution imaging of cell nucleus.^56^ These structures sacrifice nucleosomal DNA by unwrapping to preserve close contacts among neighboring nucleosomes. In addition, the presence of another 12mer promotes inter-chain interactions to stabilize extended chromatin configurations as well. Together, our results suggest that in vivo chromatin configurations can arise from the unfolding of fibril configurations as a result of tension and crowding.

## Results

### Near-atomistic modeling reproduces force-extension curve

We applied a near-atomistic model to characterize the unfolding of a 12mer chromatin with a linker length of 20 bp (Figure 1A). One bead per amino acid and three sites per nucleotide were employed to describe protein and DNA molecules, respectively, leading to a system of 23590 coarse-grained beads in size. Interactions among the coarse-grained beads were parameterized by accounting for solvent effect implicitly with physically motivated potentials (see Methods for model details). Similar approaches have been extensively used to characterize single nucleosomes^57–59^ and nucleosome oligomers^33,60^ with great success.

**Figure 1:**
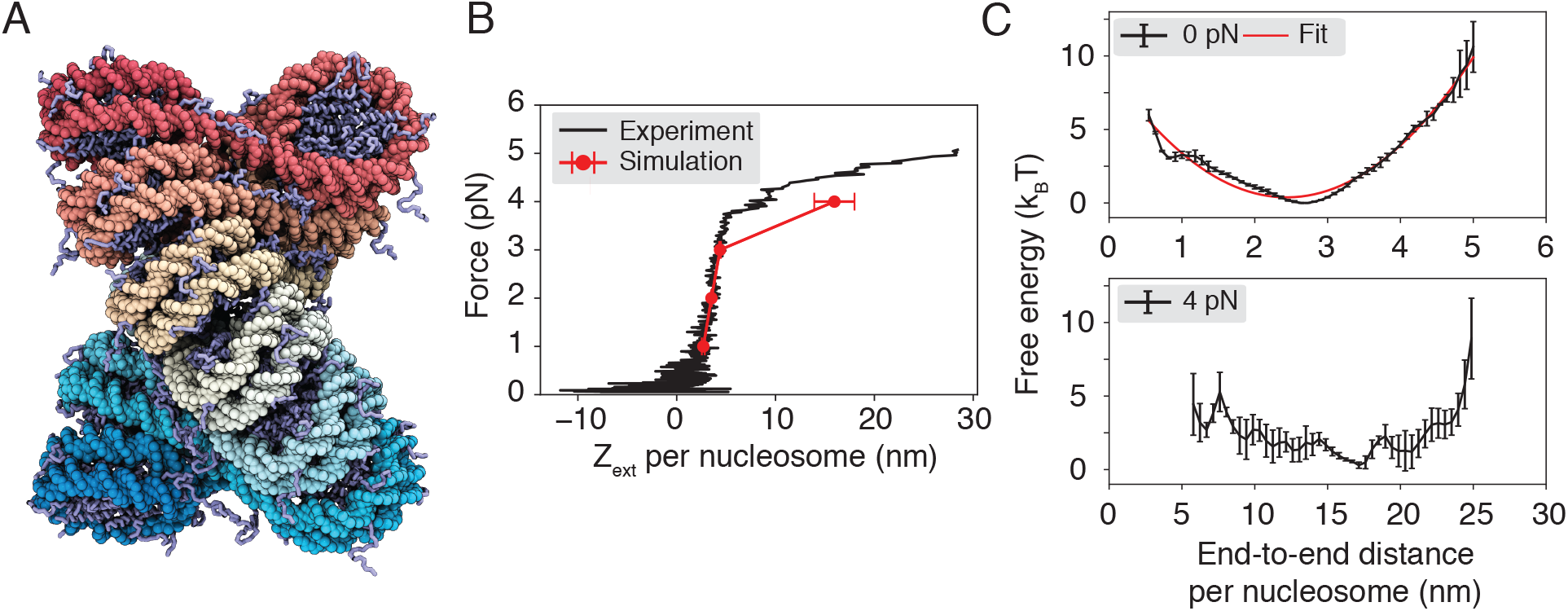
Near-atomistic modeling reproduces the force-extension curve for chromatin. (A) Illustration of the two-helix fibril chromatin structure with a linker length of 20 bp. The DNA molecule varies from red to cyan across the two ends, and histone proteins are drawn in ice blue. (B) Comparison between the simulated (red) and experimental ^55^ (black) force-extension curve. (C) Free energy profiles as a function of the DNA end-to-end distance computed with the presence of 0 pN (top) and 4 pN (bottom) extension force. A harmonic fit to the 0 pN simulation result is shown in red. Errorbars correspond to standard deviation of the mean estimated via block averaging by dividing simulation trajectories into three independent blocks of equal length.

We computed the average chromatin extension length under various pulling forces along the *z*-axis for a direct comparison with results from single-molecule pulling experiments. ^55^ Comprehensive sampling of chromatin conformations can be rather challenging because of the non-specific and strong electrostatic interactions between nucleosomes that give rise to slow dynamics. To alleviate the sampling difficulty, we carried out umbrella simulations^61^ on two collective variables that quantify the degree of nucleosomal DNA unwrapping (*q*_wrap_) and nucleosome unstacking (*d*_stack_) (Figure S1). The simulations were initialized from the most probable configurations at respective umbrella centers obtained from an exhaustive sampling of a neural network model that approximates the free energy landscape of the 12mer in terms of inter-nucleosome distances (see *Methods*). This initialization protocol attempts to prepare umbrella simulations with equilibrium configurations to avoid traps of local minima.

As shown in Figure 1B, the simulation results match well with the experimental force-extension curve measured by Kaczmarczyk et al. ^55^ In particular, we observe a linear extension regime at low forces (≤ 3 pN). The sharp increase in extension at 4 pN deviates from the linear behavior, resulting in a plateau regime. We emphasize that there are no tuning parameters in the model, and we do not make assumptions regarding stacking energies. The minor deviation at 4 pN between simulation and experiment could be due to a difference in salt concentration: while experiments were performed at 100 mM monovalent salt, we carried out the simulations at 150 mM. Reducing the salt concentration to 100 mM in simulations indeed improved the agreement in the average extension length (Figure S2).

The free energy profiles as a function of the DNA end-to-end distance are consistent with the linear and plateau regimes seen in force-extension curves (Figure 1C). In particular, at 0 pN force, the free energy curve can be well approximated with a harmonic potential, which naturally produces a linear relationship between the force and extension. Consistent with a harmonic behavior near the minimum, theoretical predictions based on the free energy profile at 0 pN match well with simulation results at 1-3 pN (Figure S3). However, the free energy profile at 4 pN is strongly anharmonic. The bottom panel shows that the curve is relatively flat over a wide range of end-to-end distances. Because of the lack of energetic penalty, a slight change in pulling force can produce significant variations in the extension length, giving rise to the observed plateau regime.

### Intermediate states support nucleosome-clutch formation

The nucleosome arrangement in extended, unfolded chromatin has been the subject of numerous studies.^39,41,46,49^ The near-atomistic simulations performed here offer a unique opportunity to produce high-resolution structures with minimal assumptions. Their success in reproducing experimental observations shown in Figures 1B and S2 supports the biological relevance of the predicted structures.

We determined representative structures at various forces to better characterize chromatin unfolding under tension (Figure 2). These structures share end-to-end distances close to the mean force-dependent extension lengths. They correspond to the central configurations of the largest clusters identified by the single-linkage algorithm^62^ using root mean squared distance (RMSD) as the distance between structures. At small forces (≤ 3 pN), though chromatin extends linearly, we do not observe a uniform extension of nucleosomes along the principal fiber axis (Figures 2 and S4). The conformational change mainly occurred in the plane perpendicular to the fiber axis via a shearing motion, causing the formation of irregular, compact structures. Such structures are more kinetically accessible as they avoid complete unstacking, which could cause a significant energetic penalty as shown by Moller et al. ^57^

**Figure 2:**
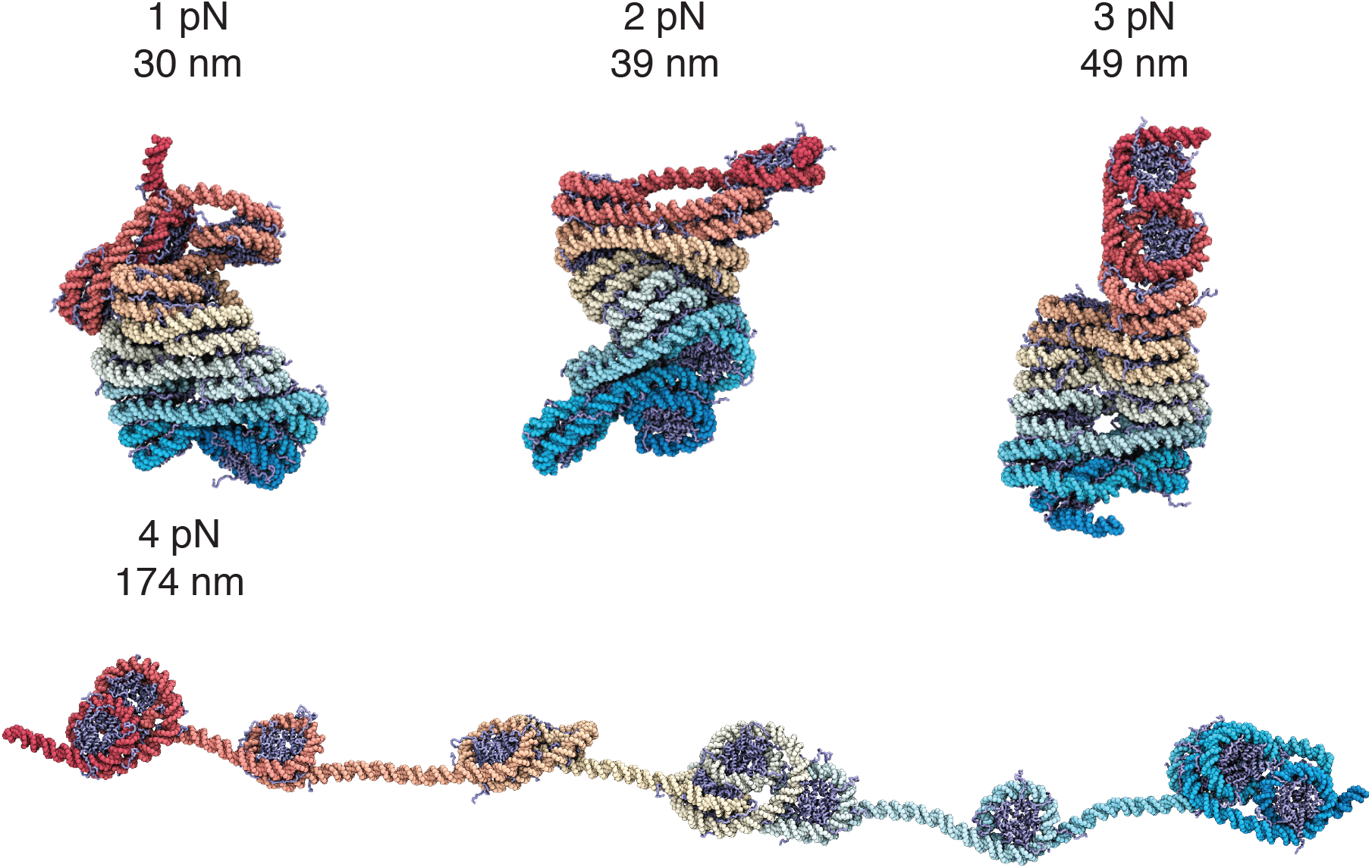
Representative chromatin structures from simulations performed under various extension forces (also see Figure S4). The values for the extension force and the end-to-end distance are provided next to the structures. The same coloring scheme as in Figure 1A is adopted here.

The preference of shearing over complete unstacking can be readily seen in Figure 3. There, we decomposed the distance between two nucleosomes into motions that are within or perpendicular to the nucleosomal plane (Figure S5). We further computed the free energy profile for the two decomposed distances under no extension force. It is evident that the energetic penalty for chromatin unfolding along the shearing direction is much smaller. Shearing can better preserve inter-nucleosome contacts as nucleosomes move away from each other, lowering the energetic penalty.

**Figure 3:**
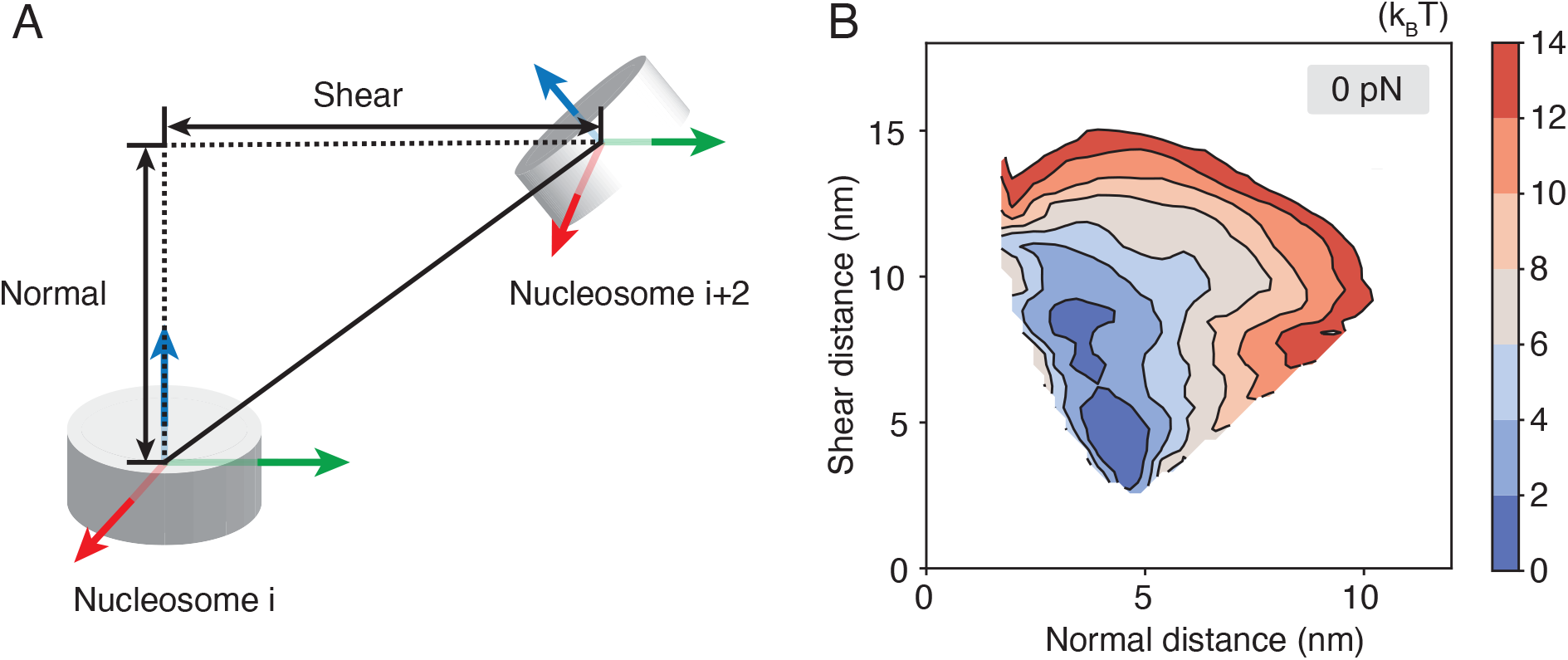
Chromatin extension favors shearing motion within the nucleosomal plane over the normal motion perpendicular to the plane. (A) Illustration of the nucleosome coordinate system and the decomposition of the inter-nucleosome distance into shearing and normal components. (B) The free energy profile as a function of the two different modes of breaking inter-nucleosome distances shown in part A.

The representative structure from 3 pN to 4 pN undergoes a dramatic transformation from a compact configuration to one with many nucleosomes losing stacking interactions. Notably, the unfolded structures fall into small clusters of nucleosomes. These structures often feature one or two nucleosomes with a highly unwrapped outer layer. Unwrapping the outer layer DNA only incurs modest energetic cost^58,63,64^ and serves as an economic strategy to extend chromatin under force. Nucleosome clutch formation is not specific to a particular end-to-end distance and can be readily seen in structures with smaller distances as well (Figure 4 and S6). We note that the nucleosomes that remain in contact are not perfectly stacked as in the crystal structure of a tetranucleosome, ^34^ but are somewhat irregular as configurations observed in prior simulations^33,52^ and in vivo experiments. ^19,35,65,66^ Further stretching the chromatin eventually leads to configurations with most of the outer nucleosomal DNA unwrapped.

**Figure 4:**
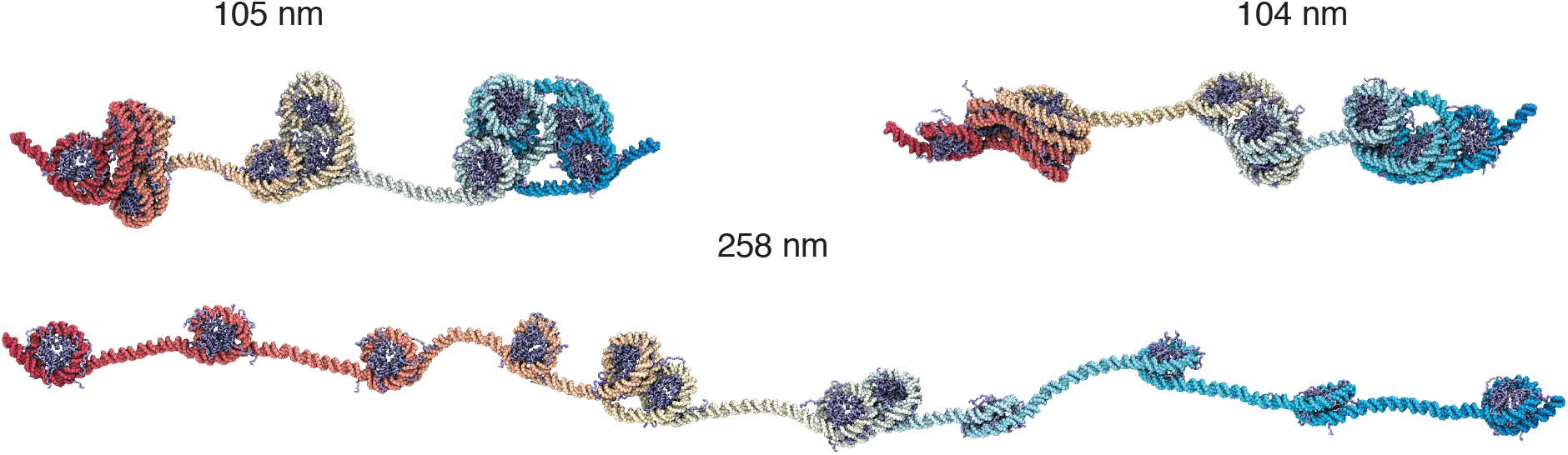
Representative chromatin structures at smaller and larger distances than the average extension at 4 pN force (see also Figure S6). These structures again support the formation of nucleosome clutches, which do not break into single nucleosomes until at very large end-to-end distances around 260 nm.

Therefore, our results suggest that chromatin unfolding is not cooperative. Nucleosomes do not uniformly unstack along the chain to extend chromatin. On the contrary, they prefer to stay in close contact as much as possible via the formation of clusters separated by unwrapped nucleosomes.

### Inter-chain contacts stabilize unfolded chromatin

The pulling simulations suggest that in vivo configurations can arise from the unfolding of chromatin fiber under tension. Inside the nucleus, chromatin is not in isolation but surrounded by other chromatin segments in a crowded environment.^20,56^ The more exposed nucleosomes in the intermediate configurations could facilitate inter-chain interactions, further stabilizing the unfolded structures.

To evaluate the impact of crowding on chromatin stability, we computed a two-dimensional free energy profile using simulations with two 12mers. The first collective variable quantifies the inter-chain contacts as the number of nucleosome pairs within 15 nm. Only pairs with one nucleosome from each chromatin segment were included to define the contacts. The other dimension measures chromatin extension using the average unstacking of the two chains, 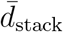. As shown in Figure 5A, configurations with close contacts between the two chromatin segments are more favorable. An representative structure for two contacting fibril chromatin identified by the single-linkage clustering algorithm is provided in Figure 5C. The contacts are mediated mainly by histone tail-DNA interactions, as can been in the inset that provides a zoomed-in view of the interface. Favorable interactions for compact chromatin are consistent with previous simulation studies that support the liquid chromatin state. ^67^

**Figure 5:**
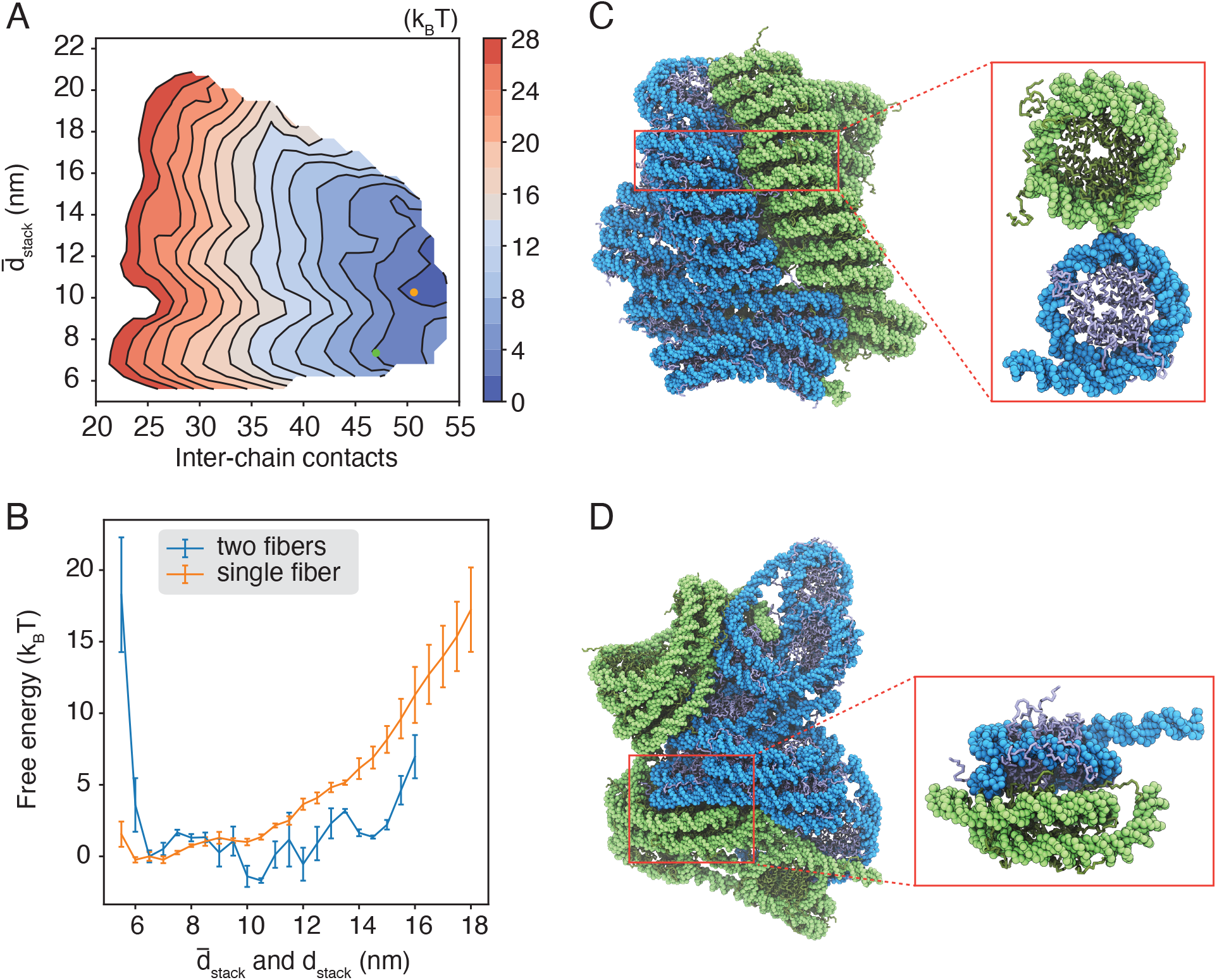
Crowding and inter-chain contacts stabilize extended chromatin configurations. (A) The free energy surface as a function of the inter-chain contacts and the average extension of the two 12mers. (B) Free energy profiles of chromatin unstacking with (blue) and without (orange) the presence of an additional 12mer. Chromatin unstacking is quantified with *d*_stack_ and 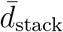 for single and two fiber simulations, respectively. (C) Representative structure for two contacting chromatin segments that maintain fibril configurations, with the corresponding collective variables indicated as the green dot in part A. The inset highlights the side-side contacts between inter-chain nucleosomes. (D) Representative structure of the free energy minimum, with the corresponding collective variables indicated as the orange dot in part A. The inset highlights the stacking interactions between inter-chain nucleosomes.

Notably, the global minimum of the free energy profile resides at larger values for 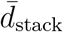 corresponding to more extended chromatin configurations. While extending chromatin is unfavorable (Figure 5B), such structures promote close contacts between nucleosomes from different chains (Figure 5D). In particular, trans-nucleosomes can now engage in stacking interactions, which are more favorable energetically compared to side-side contacts.^57^ The emergence of a new binding mode, unavailable when chromatin is constrained into fibril configurations, compensates for the energetic penalty of breaking cis-chain contacts. Further extending the chromatin leads to more intertwined structures at rather modest energetic cost (Figure S7).

### Conclusions and Discussion

We characterized the impact of tension and crowding on chromatin organization with computational modeling using a near-atomistic model. Consistent with previous studies,^48,49,53^ we observed both unwrapping of nucleosomal DNA and unstacking between nucleosomes as chromatin unfolds from the fibril configuration. However, these changes are non-uniform and are initially localized to a small set of nucleosomes, leading to the formation of nucleosome clutches separated with unwrapped nucleosomal DNA. Such intermediate structures emerge as a result of balancing intra- and inter-nucleosome interactions.

Notably, the simulated intermediate structures resemble in vivo chromatin configurations. For example, super-resolution imaging of the core histone protein H2B in interphase human fibroblast nuclei has revealed the formation of nucleosome clutches of varying size.^56^ High-resolution electron tomography studies further support the prevalence of trimers in the clutches.^19,35^ Cross-linking-based experiments that detect nucleosome contacts in situ support nucleosome clutches with tri- or tetranucleosome as well. ^65,68^ Our results generalize the findings from a previous study on tetra-nucleosomes.^33^ They support in vivo chromatin structures as folding intermediates of the fibril configuration. While the fibril structure with stacked nucleosomes is the most stable configuration for chromatin in isolation, tension can promote unfolding. The clutch configurations are mostly seen under 4 pN, a value that is indeed within the range expected from molecular motors.^21^

We further showed that the unfolded chromatin could promote inter-chain contacts, leading to the formation of interdigitated structures. Such structures present an alternative binding mode compared to the close contacts between two fibril configurations. In addition to supporting chromatin unfolding in a crowded environment, the interdigitated structures suggest that chromatin may, in fact, form gels at high density inside the nucleus. Gelation can form due to the stacking interactions between exposed nucleosomes from different chains, which are stronger than side-side interactions that are only accessible for nucleosomes in closely stacked fibers. Therefore, the two binding modes could help understand the observation of both liquid and gel state of chromatin mixtures.^67,69–72^

In addition to tension and crowding, additional factors could impact chromatin organization as well, including histone modifications,^73–75^ linker DNA length,^12,60,76–78^ chromatin regulators,^40,69,79–81^ and salt concentrations.^66,82–84^ The near-atomistic simulations can be generalized straightforwardly to investigate the role of many factors on the formation of nucleosome clutches. One possible limitation of the modeling approach here is our use of implicit solvation. Additional force field development would be needed to quantitatively predict the impact of salt concentration and multivalent ions on chromatin organization and inter-chain contacts.

## Methods

### Near-atomistic modeling of chromatin organization

We applied a near-atomistic model to study a chromatin segment with twelve nucleosomes. The structure-based model ^85,86^ was used to represent protein molecules with one bead per amino acid and stabilize the tertiary structure of the histone octamer while maintaining the conformational flexibility of disordered tail regions. Protein molecules from different nucleosomes interact through both an electrostatic and amino acid-specific potential.^87^ We represented the DNA molecule with three beads per nucleotide using the 3SPN model.^88^ Protein-DNA interactions were described with the screened Debye-Hückel potential at a salt concentration of 150 mM and the Lennard-Jones potential for excluded volume effect. The near-atomistic model has been used extensively in prior studies to investigate protein-protein/protein-DNA interactions,^81,89^ the energetics of single nucleosome unwinding, ^58,63^ nucleosome-nucleosome interactions,^57^ and the folding pathways of a tetra-nucleosome.^33^ More details on the model setup and force field parameters can be found in the Supporting Information.

The software package LAMMPS^90^ was used to perform molecular dynamics simulations with periodic boundary conditions and a time step of 10 fs. The length of the cubic simulation box was set as 2000 nm. We used the Nosé-Hoover style algorithm^91^ to maintain the temperature at 300 K with a damping constant of 1 ps. The globular domains of histone proteins and the inner layer of nucleosomal DNA were rigidified, while leaving the outer layer, linker DNA, and disordered histone tails flexible. As shown in Ref. 33, this treatment does not impact the accuracy in sampling inter-nucleosome interactions but significantly reduces the computational cost.

Most simulations lasted for more than 10 million steps. Exact trajectory lengths are provided in Tables S1. We computed the error bars by dividing the data into three equal-length, non-overlapping blocks and calculated the respective quantities using data from each block. The standard deviations of the three estimations were used to estimate the error of the mean.

### Force extension curves from enhanced sampling

To characterize chromatin structures under tension and compute force-extension curves, we introduced two collective variables that monitor the important degrees of freedom for chromatin unfolding. The first variable, *d*_stack_, measures the average center of mass distance between the *i*-th and (*i* + 2)-th nucleosomes. For small values of *d*_stack_, nucleosomes are stacked on top of each as in the zigzag conformation. ^13,34^ The second variable, *q*_wrap_, quantifies the average degree of nucleosome unwrapping. The two variables can better differentiate the various chromatin conformations and capture the energetic cost of extension than the DNA end-to-end distance. Mathematical expressions for the two variables are provided in the SI. For extension forces less than 4 pN, we carried out a set of two-dimensional umbrella sampling based on *q*_wrap_ and *d*_stack_. *q*_wrap_ was restricted to centers from 0.45 to 0.90 with a spacing of 0.15 and a spring constant of 50.0 kcal/mol. *d*_stack_ was restricted to centers from 10.0 nm to 30.0 nm with a spacing of 5 nm and a spring constant of 0.05 kcal/mol/nm^2^. Additional simulations were added to further improve the overlap between umbrella windows and the convergence of free energy calculations. At 4 pN, chromatin can adopt fully unstacked structures with large end-to-end distances. Covering the entire accessible phase space with two dimensional umbrella simulations becomes too costly computationally. Therefore, we restricted to one-dimensional free energy calculations using *d*_stack_ as the collective variable. All simulations were initialized from the most probable configurations predicted by a neural network model for chromatin stability under the same umbrella biases (see below for details). Details of the umbrella centers and spring constants used in all simulations are provided in Table S1.

### Facilitating conformational sampling with a neural network model for chromatin

Conformational sampling for the 12mer is challenging because of the many possible degenerate configurations. For example, both unstacking and unwrapping can extend chromatin, and different combinations of the two from various nucleosomes can result in a large number of structures that share similar end-to-end distances. Conformational transitions among them are slow due to considerable energetic barriers arising from non-specific electrostatic interactions.

To alleviate the sampling problem, we introduced a neural network model for the 12mer. As detailed in the SI, the model quantifies the stability and the free energy of chromatin structures using inter-nucleosome distances. It was parameterized using mean forces estimated with near-atomistic simulations for 10,000 independent tetra-nucleosome configurations. The neural network model is computationally efficient and allows exhaustive Monte Carlo sampling to determine the most likely chromatin structures at a given setup. These structures were provided to initialize near-atomistic simulations and free energy calculations. The neural network model is not perfect due to approximations introduced when building the free energy surface with tetra-nucleosome calculations. However, it does reproduce the force-extension curve reasonably well at the lower force regime (Figure S8). We only used the neural network model for conformational exploration, and all quantitative results presented in the manuscript were obtained with near-atomistic simulations.

### Exploring the impact of crowding on chromatin extension

To study the impact of crowding on chromatin organization, we computed the free energy profile as a function of two collective variables that measure intra- and inter-chain contacts. Umbrella sampling was used to enhance conformational exploration, and details on the restraining centers and constants are provided in Table S2.

Umbrella simulations were initialized from configurations in which the two chains were separated far apart from each other with zero contacts. For simulations biased toward small values of 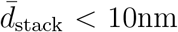, we prepared each chromatin with a two-helix zigzag configuration that resembles the cryo-EM structure^13^ (see Figure 1A). The rest of simulations were initialized with extended chromatin configurations predicted by the neural network model. More simulation details can be found in the SI.

## Acknowledgement

This work was supported by the National Institutes of Health (Grant R35GM133580) and the National Science Foundation (Grant MCB-2042362). We thank Dr. van Noort for sharing the experimental force-extension data.

## Author contributions

Conceptualization: SML, XCL, BZ

Methodology: SML, XCL, BZ

Investigation: SML, XCL, BZ

Visualization: SML, XCL, BZ

Supervision: BZ

Writing original draft: SML, XCL, BZ

Writing review & editing: SML, XCL, BZ

## Data and materials availability

Data presented in this study is available upon reasonable request to the corresponding author.

