## Supporting Information for "Chromatin fiber breaks into clutches under tension and crowding"

#### Summary

**Details of Near-atomistic Simulations** **S-2**

System Setup . . . . . . . . . . . . . . . . . . . . . . . . . . . . . . . . . . . . . S-2

Force Field Setup . . . . . . . . . . . . . . . . . . . . . . . . . . . . . . . . . . . S-3

Free Energy Profiles for Chromatin Under Tension . . . . . . . . . . . . . . . . . S-4

Initial configurations from the neural network model . . . . . . . . . . . . . . . S-6

One-dimensional free energy calculations at 4 pN . . . . . . . . . . . . . . . . S-7

Estimating the extension per nucleosome from experimental data . . . . . . S-7

Theoretical predictions of chromatin extension along the $z$ axis . . . . . . . . S-8

Decomposing Inter-nucleosome distances into Shear and Normal Motion . . . . . . S-9

Free Energy Calculations for Two Interacting 12mers . . . . . . . . . . . . . . . . S-10

**Neural Network Model for the 12mer Chromatin** **S-11**

Numerical simulations of the Neural Network Model . . . . . . . . . . . . . . . . . S-12

### Details of Near-atomistic Simulations

We carried out all molecular dynamics simulations with the software LAMMPS.[S1] Umbrella sampling was performed using collective variables implemented by the Plumed package.[S2] We applied the weighted histogram method[S3,S4] and fastmbar[S5] to process the data and computed the unbiased extension length at a given force.

#### System Setup

We built a structural model for the chromatin with 12 nucleosomes and 20-bp linker DNA following two steps. We first connected 12 individual nucleosomes into a continuous segment without much regard to the overall chromatin topology. We then aligned the DNA model to a template that closely resembles the cryo-EM structure with a two-start fibril organization.[S6]

We connected individual nucleosomes to build a 12mer chromatin as follows. The nucleosome unit with 167-bp of DNA was extracted from the tetranucleosome X-ray structure (PDB ID:1ZBB).[S7] The DNA was taken as residues 158-324 of chain I and the corresponding complementary segment from chain J. There are no extra base pairs at the entry side of the nucleosome in this setup, but 20-bp linker DNA exists at the exiting end. The resulting sequence with the core DNA (147bp) in underline is

ACAGGATGTAACCTGCAGATACTACCAAAAGTGTATTTGGAAACTGCTCCAT
CAAAAGGCATGTTCAGCTGGATTCCAGCTGAACATGCCTTTTGATGGAGCAG
TTTCCAAATACACTTTTGGTAGTATCTGCAGGTGATTCTCCAGGGCGGCCAG
TACTTACATGC

We further replaced the coordinates for histone proteins with that from PDB ID: 1KX5,[S8] which resolved the coordinates for histone tails.

We added one additional DNA base pair at the end of the linker DNA as sticky ends using the software 3DNA[S9] for alignment between neighboring nucleosomes. This 168-bp segment is the building block for constructing the dodecamer. For example, to extend chromatin with $n$ nucleosomes, we align the 168-th bp of the $n$-th nucleosome with the first bp of the $(n+1)$-th. The alignment determines the orientation of the $(n+1)$-th nucleosome, and the fiber is extended by one after removing the overlapping nucleotides. For the last (12-th) nucleosome, we deleted the linker DNA to build the dodecamer with 1984 bp of DNA. The resulting all-atom model was coarse-grained into the near-atomistic model with in-house scripts.

While the above procedure succeeds at building an all-atom model for the 12mer, the precise topology of the resulting structure cannot be controlled easily. To construct a two-start fibril configuration resembling the compact and twisted Cryo-EM structure,[S6] we aligned the model to a two-start fiber structure built by the software fiberModel, as detailed below. The structural alignment was performed using MDAnalysis[S10,S11] with RMSD coordinate fitting.[S12,S13]

The template was generated by the software fiberModel as the lowest energy configuration.[S14] FiberModel optimized fiber configurations by utilizing a series of geometric parameters, including the height per nucleosome along the fiber axis ($h$), the rotation angle per nucleosome around the fiber axis ($\theta$), the radius of the fiber ($R$), and three Euler angles that determine the direction of each nucleosome $(\alpha, \beta, \gamma)$. To build a fibril chromatin structure, we set initial values for the parameters $(h, \theta, R, \alpha, \beta, \gamma)$ as (2.34 nm, 2.88, 7.3 nm, -3.14, 0.622, 0) as estimated from the cryo-EM structure.[S6] Also, each nucleosomes was treated as a cylinder with the radius and height of 5.2 and 4.5 nm. We then used fiberModel to optimize the chromatin structure based on parameters $\alpha$ and $\gamma$, while keeping the other parameters fixed. The optimization utilized the basin hopping global search technique.[S15] The final strucutre aligned with FiberModel template is shown in Figure 1A.

#### Force Field Setup

We used the same force fields as in the tetra-nucleosome study[S16] to simulate the 12mer. The 3SPN.2C DNA model[S17] was adopted to model each nucleotide with three coarse-grained beads for phosphate, sugar, and base, respectively. The C$\alpha$ structure-based model[S18] was adopted to simulate the conformational dynamics of individual histone proteins. Both bonded and nonbonded interactions were generated based on the nucleosome crystal structure (PDB ID: 1KX5). For nonbonded contact potentials, two residues were considered in contact when their minimum distance is smaller than 6Å, implemented using the Shadow algorithm.[S19] We further scaled the default interaction strength[S4] by 2.5 to prevent proteins from unfolding at 300K. To model the disordered portions of the histones, we removed the dihedral and contact potentials for disordered residues not included in the core histones (residue ID: 44-135, 160-237, 258-352, 401-487, 531-622, 647-724, 745-839, 888-974). The IDs continuously index residues from chain A to chain H of the crystal structure with PDB ID: 1KX5.

In addition, residue-specific protein-protein interactions were introduced with the Miyazwa-Jernigan (MJ) potential[S20] and scaled by a factor of 0.4. In a previous study, we showed that the scaled MJ potential provides a balanced modeling of the radius of gyration for both folded and disordered proteins.[S16]

Protein-DNA interactions include the electrostatic potential modeled at the Debye-Hückel level with a salt concentration of 150mM. In addition, a weak, non-specific Lennard-Jones potential was applied between all protein-DNA beads. Detailed expression for these potentials can be found in Ref. S21.

#### Free Energy Profiles for Chromatin Under Tension

We defined two collective variables to explore chromatin configurations and compute free energy profiles. The unwrapping variable, $q_{\text{wrap}}$, quantifies DNA unwrapping using the distance between neighboring nucleosome $d_{i,i+1}$. It is defined as

$$q_{\text{wrap}} = \frac{1}{11} \sum_{i=1}^{11} \exp \left[ -\frac{(\max(d_{i,i+1}, d_o) - d_o)^2}{2\sigma_w^2} \right]. \tag{S1}$$

$d_o = 15$ nm is close to the distance between two neighboring nucleosomes in the PDB structure (PDB ID: 1KX5),[S8] and we used $\sigma_w = 4$ nm. The function max selects the larger value of the two distances. The above definition makes use of the geometric constraint that increase in the distances between neighboring nucleosomes ($d_{i,i+1}$) can only arise from nucleosome unwrapping. The unstacking variable, $d_{\text{stack}}$, measures the mean distance between $i$-th and $(i+2)$-th nucleosomes as

$$d_{\text{stack}} = \frac{1}{10}\sum_{i=1}^{10} d_{i,i+2}. \tag{S2}$$

Umbrella simulations with the two collective variables at forces (0-3 pN) were carried out to compute the free energy profiles. To compare our simulations with force-extension experiments, we applied force $f_{\text{ext}}$ along the $z$-axis projection of the DNA end-to-end distance ($L_z$). The two DNA ends were defined as the geometric centers of all the coarse-grained beads for the first and last five base pairs. The potential energy function of these simulations at center $(q_o, d_o)$ and force $f_{\text{ext}}$ is defined as

$$U_{\text{biased}} = U(\mathbf{r}) + \frac{\kappa_q}{2}(q_{\text{wrap}} - q_o)^2 + \frac{\kappa_d}{2}(d_{\text{stack}} - d_o)^2 - f_{\text{ext}} L_z, \tag{S3}$$

where $U(\mathbf{r})$ corresponds to the interaction energy defined by the force field. The umbrella centers $(q_o, d_o)$ were initially placed on a uniform grid $[0.45 : 0.15 : 0.9] \times [10 : 5 : 30]$. We introduced additional centers to improve the overlap between umbrella simulations. A complete list of the umbrella centers and the restraining constants is provided in Table S1.

At the extension force of 4 pN, the 12mer can adopt configurations that cover a wide range of $q_{\text{wrap}}$ and $d_{\text{stack}}$. Uniform sampling of the entire accessible phase space becomes too costly computationally. Therefore, we only carried out one-dimensional umbrella simulations with $d_{\text{stack}}$ as the collective variable.

**Initial configurations from the neural network model**

Conformational sampling of the near-atomistic model is challenging due to strong but non-specific electrostatic interactions. We initialized the umbrella sampling simulations with the most probable configurations predicted by a neural network under a similar setup to alleviate the sampling challenge. As detailed in the *Section: Neural Network Model for the 12mer Chromatin*, the neural network model quantifies the stability of chromatin configurations using inter-nucleosome distances. It's computationally efficient and allows equilibrium sampling of chromatin configurations.

For a near-atomistic umbrella simulation centered at $(q_o^N, d_o^N)$ with extension force $f_{\text{ext}}$, we carried out replica exchange Monte Carlo sampling of the following biased free energy

$$F_{\text{biased}} = F(\mathbf{d}) + \frac{\kappa_q}{2}(q_{\text{wrap}} - q_o^N)^2 + \frac{\kappa_d}{2}(d_{\text{stack}} - d_o^N)^2 - f_{\text{ext}}L, \tag{S4}$$

with $\kappa_q = 47.8$ kcal/mol and $\kappa_d = 4.78 \times 10^{-2}$ kcal/(mol $\cdot$ nm$^2$). $\mathbf{d}$ corresponds to the list of inter-nucleosome distances, and $L$ is the distance between the first and the last nucleosomes. See *Section: Monte Carlo Sampling of the Machine Learning Model* for sampling details. We used the samples collected in the final 300000 steps of the 300K replica for a K-means clustering analysis with 10 centers. The configuration closest to the center of the largest cluster was selected as the most probable configuration.

The neural network model represents chromatin structures with inter-nucleosome distances. We performed short targeted molecular dynamics simulations starting from the two-helix fiber to build near-atomistic model structures consistent with the most probable configurations from the neural network sampling. These simulations bias on all the inter-nucleosome distances with a restraining constant of 23.9 kcal/(mol $\cdot$ nm$^2$) for approximately 300000 steps. The end configurations of these simulations were used to initialize the near-atomistic umbrella simulations.

#### One-dimensional free energy calculations at 4 pN

For 4 pN extension force, we carried out umbrella simulations with the energy function

$$U_{\text{biased}} = U(\mathbf{r}) + \frac{\kappa_d}{2}(d_{\text{stack}} - d_o)^2 - f_{\text{ext}} L_z, \tag{S5}$$

where $\kappa_d = 0.05$ kcal/(mol · nm$^2$). We chose $d_o$ from 10 to 50 nm with a step size of 2.5 nm.

Two sets of simulations with different initial configurations were performed at each umbrella window. The initial configurations were constructed using the most probable structures predicted by the neural network model. We performed two-dimensional umbrella bias simulations of the neural network model as detailed in *Section: Initial configuration from neural network model.* Two sets of most probable structures were obtained from umbrella simulations with bias centers at $q_o^N = 0.45$ and 0.60, as well as $d_o^N = 10.0$ nm, 15.0 nm, 20.0 nm, 25.0 nm, and 30.0 nm. We used the neural network configurations from simulations with $d_o^N$ values closet to $d_o$ to initialize the corresponding near-atomistic simulations.

#### Estimating the extension per nucleosome from experimental data

We processed the force-extension curve from single-molecule force spectroscopy experiments to compute the extension per nucleosome as follows. The extension length from experiments includes contributions from both the DNA handle and the chromatin. Following previous study,[S22] we estimate the DNA handle extension as

$$L_{z,\text{handle}} = L_{\text{c,handle}} \times \left(1 - \frac{1}{2}\sqrt{\frac{k_B T}{f_{\text{ext}} A}} + \frac{f_{\text{ext}}}{S}\right) \tag{S6}$$

where $k_B$ is Boltzmann constant, $T$ is temperature, $A$ is the persistence length of DNA, $f_{\text{ext}}$ is the extension force along the $z$ axis, and $S$ is the stretching modulus. We used $A = 50$ nm, $S = 900$ pN, and $T = 300$ K. The contour length of the DNA handle, $L_{c,\text{handle}}$, is estimated as

$$L_{c,\mathrm{handle}} = [n_{\mathrm{bp}} - \mathrm{NRL} \times (n_{\mathrm{nucl}} - 1) - 147]b \tag{S7}$$

where $n_{\mathrm{bp}}$ is the total number of base pairs in DNA, NRL is nucleosomal repeat length, $n_{\mathrm{nucl}}$ is the total number of nucleosomes, and $b$ is the length of each base pair. We used NRL = 167 bp, $n_{\mathrm{nucl}} = 25$, $n_{\mathrm{bp}} = 7045$ bp, and $b = 0.34$ nm.

Subtracting the extension of the DNA handle from the total extension length $L_z$, the extension per nucleosome can be estimated as

$$L_{z,\mathrm{nucl}} = \frac{L_z - L_{z,\mathrm{handle}}}{n_{\mathrm{nucl}} - 1}. \tag{S8}$$

#### Theoretical predictions of chromatin extension along the $z$ axis

To better understand the linear extension of chromatin at small forces, we introduced an analytical model based on simulation results without extension force.

We approximate the unbiased free energy profile for chromatin extension at zero force with a harmonic function, $F(L) = a(L-L_0)^2+b$, where $L$ is the extension length, i.e., the end-to-end distance. The parameters were obtained by a least-squares fitting to the simulation data presented in Figure 1C of the main text, resulting in $a = 1.200 \times 10^{-2}\ k_BT/\mathrm{nm}^2$, $L_0 = 26.83$ nm, and $b = 0.3820\ k_BT$. The corresponding free energy profile with an extension force $f$ along the $z$-axis can be defined as $F_f(L) = F(L) - fL\cos\theta$, where $\theta$ is the azimuthal angle (i.e. the angle between the fiber axis and the $z$-axis). From this expression, the average extension along the $z$-axis can be computed as

$$\langle L_z \rangle_f = \frac{\int_0^\infty \int_0^\pi L\cos\theta e^{-\beta F_f(L)} L^2 dL \sin\theta d\theta}{\int_0^\infty \int_0^\pi e^{-\beta F_f(L)} L^2 dL \sin\theta d\theta} \tag{S9}$$

Numerical integration of the above equation led to $\langle L_z \rangle_f = 0$, 33.87, 45.76, and 56.36 nm for extension force of 0, 1, 2, and 3 pN, respectively. The extension per nucleosome along the $z$ axis ($Z_{\text{ext}}$ per nucleosome) is defined as $\langle L_z \rangle_f/11$ and shown in Figure S3.

#### Decomposing Inter-nucleosome distances into Shear and Normal Motion

As discussed in the main text, two distinct motions can increase the distance between $i$-th and $(i+2)$-th nucleosomes and the collective variable $d_{\text{stack}}$. To quantitatively characterize these two motions, we introduced a coordinate system for each nucleosome. Following de Pablo and coworkers,[S23] we defined the origin of the coordinate system using the geometric center of residues 63-120, 165-217, 263-324, 398-462, 550-607, 652-704, 750-811, and 885-949. The IDs continuously index residues from chain A to chain H of PDB 1KX5. Two additional points were introduced to define the nucleosomal plane using the geometric center of the dyad that includes CG atoms 81-131, 568-618, and the geometric center of CG atoms 63-120, 165-217, 750-811, and 885-949. Two unit vectors, $\mathbf{u}$ and $\mathbf{v}$, can then be defined using the vectors pointing from the origin to the dyad and the third point. Atoms in the third point were chosen such that $\mathbf{u}$ and $\mathbf{v}$ are approximately orthogonal to each other. The unit normal vector $\mathbf{w}$ for nucleosome plane can then be defined as the cross product, $\mathbf{u} \times \mathbf{v}$. An illustration of the various axes is provided in Figure S5.

With the nucleosomal axes defined above, the distances between two nucleosomes can be decomposed to the distances within the nucleosomal plane, i.e., shearing, and the distance perpendicular to the plane, i.e., unstacking. Denoting the vector from nucleosome $i$ to nucleosome $i+2$ as $\mathbf{d}_{i,i+2}$ (here we use the distance between the coordinate origins for the two nucleosomes), the corresponding normal and shear distances are $d^n_{i,i+2} = |\mathbf{d}_{i,i+2} \cdot \mathbf{w}_i|$ and $d^s_{i,i+2} = \sqrt{|\mathbf{d}_{i,i+2}|^2 - {d^n_{i,i+2}}^2}$. The normal and shear distances for the 12mer chromatin, $d_n$ and $d_s$, are defined using the mean values of all nucleosome $i$ and $i+2$ pairs as $d_n = \frac{1}{10}\sum_{i=1}^{10} d^n_{i,i+2}$ and $d_s = \frac{1}{10}\sum_{i=1}^{10} d^s_{i,i+2}$.

#### Free Energy Calculations for Two Interacting 12mers

To quantify the impact of chromatin-chromatin interactions and crowding on the stability of fibril configurations, we carried out simulations with two chromatin segments. Umbrella sampling was performed using two collective variables. The first variable quantifies the average extension of the two 12mers with $\bar{d}_{\text{stack}}$ defined as

$$\bar{d}_{\text{stack}} = \frac{1}{2}(d^1_{\text{stack}} + d^2_{\text{stack}}). \tag{S10}$$

$d_{\text{stack}}$ is defined in Eq. S2 and $1, 2$ indecies the two 12mers. The second variable measures the number of contacts between the two chromatins. Contacts were defined at the nucleosome level, and a pair of nucleosomes is denoted as in-contact if the distance between their geometric centers ($d_{i,j}$) is less than 15 nm. Mathematically, the interchain contacts, $C$, is defined as

$$C = \sum_{i=1}^{12} \sum_{j=13}^{24} \frac{1 - (d_{i,j}/d_o)^6}{1 - (d_{i,j}/d_o)^{12}}, \tag{S11}$$

where $i$ and $j$ indices over nucleosomes from the two 12mers and $d_o = 15$ nm.

We biased the simulations towards various collective variable values for a comprehensive exploration of the phase space. Details of the umbrella centers and force restraints used in our simulations are provided in Table S2. Simulations with umbrella centers $\bar{d}_{\text{stack}}$ biased to values $\leq$ 10nm were initialized using the two-helix fibril configuration for each chromatin placed at $\geq$ 20 nm apart. The rest of the simulations were initialized with extended chromatin configurations extracted from the neural network simulations. Chromatin configurations in simulations with 4 pN extension force at umbrella centers $d_o$ were adopted here for simulations that biased $\bar{d}_{\text{stack}}$ to the same values. Only chromatin configurations for the first set of simulations with 4 pN extension force were used here (see *Section: One-dimensional free energy calculations at 4 pN*). To ensure the equilibrium of the system, the initial five million steps of each of the simulation trajectories were discarded in our free energy calculations.

#### Neural Network Model for the 12mer Chromatin

To facilitate conformational sampling of the 12mer, we introduced a neural network model to quantify the free energy of chromatin configurations as a function of inter-nucleosomal distances.

The neural network model is a generalization of the free energy surface for a tetra-nucleosome determined in a previous study.[S16] The previous study provided a neural network model that can compute the free energy of a tetra-nucleosome given all the distances between any two nucleosomes, and we aimed to generalize this model thus we could estimate the free energy of 12mer. We defined $F_n(1\ldots,n)$ as the free energy of an oligomer including $n$ nucleosomes of indices $1,\ldots,n$. We assumed that each nucleosome with index $i$ could only interact with nucleosomes $i \pm 1/2/3$, while ignoring nucleosome pair interactions beyond tetramers. Under this assumption, the free energy of $n+1$ nucleosomes ($F_{n+1}$) can be determined from the following recursive relationship as

$$F_{n+1}(1,\ldots,n+1) = F_n(1,\ldots,n) + F_4(n-2,n-1,n,n+1) - F_3(n-2,n-1,n). \quad \text{(S12)}$$

Subtracting the free energy ($F_3$) avoids the double-counting from adding the tetrameric contribution ($F_4$).

The trimer free energy is estimated as follows. Assuming that the fourth nucleosome is far away from the rest of the three and its interaction with them can be ignored, the free energy difference between the two should be a constant. Therefore, we have

$$F_3(d_{1,2},d_{1,3},d_{2,3}) = F_4(d_{1,2},d_{1,3},d_{2,3},d_{1,4}=d_{2,4}=d_{3,4}=15\text{nm}) + \text{const.} \quad \text{(S13)}$$

Here $d_{i,j}$ refers the distance between nucleosome $i$ and $j$. The distances from the fourth nucleosome to the other three (i.e. $d_{1,4}, d_{2,4}, d_{3,4}$) were set as 15 nm. For $d_{3,4}$, this value is comparable to the distance between neighboring nucleosomes in the PDB structure for a tetra-nucleosome to avoid significant DNA unwrapping or DNA overstretching. It is also large enough to unstack $i$ and $i \pm 2$ nucleosomes and to dissociate $i$ and $i \pm 3$ nucleosome contacts, based on previous computational results.[S23] The effectiveness of this generalized neural network model is verified based on the fact that it can accurately predict the extension at different forces (Figure S8).

Given all the $d_{i,i\pm1/2/3}$ and assuming a left-handed helix, the relative position of each nucleosome can be uniquely determined geometrically, as long as the distances satisfy some geometric requirements such as triangle inequality. After determining the relative position of each nucleosome, the full set of distances (i.e. distances between any two different nucleosomes) was used to bias near-atomistic simulations towards the most probable configurations predicted by the neural network sampling.

#### Numerical simulations of the Neural Network Model

We used the replica-exchange Monte Carlo algorithm to explore the free energy surface defined by the neural network. 20 Replicas with temperatures as geometric sequence from 300 K to 2000 K were used. 500000 steps of simulations were performed for each replica. The initial 20000 steps were used to optimize the MC simulation step size so that the mean acceptance rate of MC movement is $\sim 0.20 - 0.25$. The exchange between two neighboring replicas was attempted every 50 steps. We used the samples collected in the final 300000 steps of the replica at 300 K for analysis.

Table S1: Summary of umbrella simulation details for free energy calculations at various extension forces. The format for umbrella centers, "start:end:step", indicates the a series of values from "start" to "end" with a spacing of "step". The two restraining constants are shown in the format "($\kappa_{q_{\mathrm{wrap}}}$ (kcal/mol), $\kappa_{d_{\mathrm{stack}}}$ (kcal/(mol · nm$^2$)))".

| Extension force (pN) | Umbrella center: $q_{\mathrm{wrap}}$ | Umbrella center: $d_{\mathrm{stack}}$ (nm) | Restraining constants | Simulation length (million steps) |
|---|---|---|---|---|
| 0 | 0.45:0.90:0.15 | 10.0:30.0:5.0 | (50, 0.05) | 10.5 |
| 0 | 1.00 | 6.0:10.0:0.5 | (47.8, 1.20) | 10 |
| 0 | 1.00 | 10.0:15.0:2.5 | (47.8, 0.120) | 10 |
| 0 | 1.00 | 12.5:15.0:2.5 | (47.8, 0.478) | 10 |
| 0 | 0.75:0.95:0.05 | 6.0:10.0:0.5 | (120, 1.20) | 10 |
| 0 | 0.90:0.95:0.05 | 10.0:15.0:2.5 | (120, 0.120) | 10 |
| 0 | 0.90:0.95:0.05 | 12.5:15.0:2.5 | (120, 0.478) | 10 |
| 0 | 0.80:0.85:0.05 | 10.0:20.0:2.5 | (120, 0.0120) | 10 |
| 0 | 0.75:0.85:0.05 | 12.5:20.0:2.5 | (120, 0.478) | 10 |
| 1 | 0.45:0.90:0.15 | 10.0:30.0:5.0 | (50, 0.05) | 10 |
| 2 | 0.45:0.90:0.15 | 10.0:30.0:5.0 | (50, 0.05) | 10 |
| 3 | 0.45:0.90:0.15 | 10.0:30.0:5.0 | (50, 0.05) | 15 |
| 3 | 0.45 | 10.0:20.0:5.0 | (50, 0.2) | 15 |
| 3 | 0.60 | 10.0:20.0:5.0 | (50, 0.2) | 15 |
| 3 | 0.75 | 10.0:30.0:5.0 | (50, 0.2) | 15 |
| 3 | 0.90 | 10.0:30.0:5.0 | (50, 0.2) | 15 |
| 4 (1st set) | n.a. | 10.0:30.0:2.5 | (0, 0.05) | 24.5 |
| 4 (2nd set) | n.a. | 10.0:30.0:2.5 | (0, 0.05) | 25 |

Table S2: Summary of umbrella simulation details for free energy calculations with two 12-mers. The same format as in Table S1 is adopted here, and the units for the two restraining constants are $\kappa_C$(kcal/mol), $\kappa_{\bar{d}}$(kcal/(mol $\cdot$ nm$^2$).

| Umbrella center: $C$ | Umbrella center: $\bar{d}$ (nm) | Restraining constants | Simulation length (million steps) |
|---|---|---|---|
| 30.0:45.0:5.0 | 10.0:25.0:2.5 | (0.1, 0.05) | 20 |
| 10.0:20.0:5.0 | 6.0:10.0:0.5 | (0.120, 1.20) | 10 |
| 10.0:20.0:5.0 | 10.0:25.0:2.5 | (0.478, 0.239) | 10 |
| 10.0:20.0:5.0 | 10.0:25.0:2.5 | (0.120, 0.0478) | 10 |
| 10.0:20.0:5.0 | 9.0:9.5:0.5 | (0.120, 4.78) | 10 |
| 25.0:45.0:5.0 | 6.0:10.0:0.5 | (0.120, 1.20) | 20 |
| 25.0 | 10.0:25.0:2.5 | (0.120, 0.0478) | 20 |
| 25.0:45.0:5.0 | 9.0:9.5:0.5 | (0.120, 4.78) | 20 |
| 25.0 | 10.0 | (0.120, 4.78) | 20 |
| 25.0:45.0:5.0 | 10.0 | (0.478, 0.239) | 20 |
| 30.0:45.0:5.0 | 10.0 | (0.120, 0.0478) | 20 |
| 30.0:40.0:5.0 | 12.5:15.0:2.5 | (0.478, 0.239) | 20 |

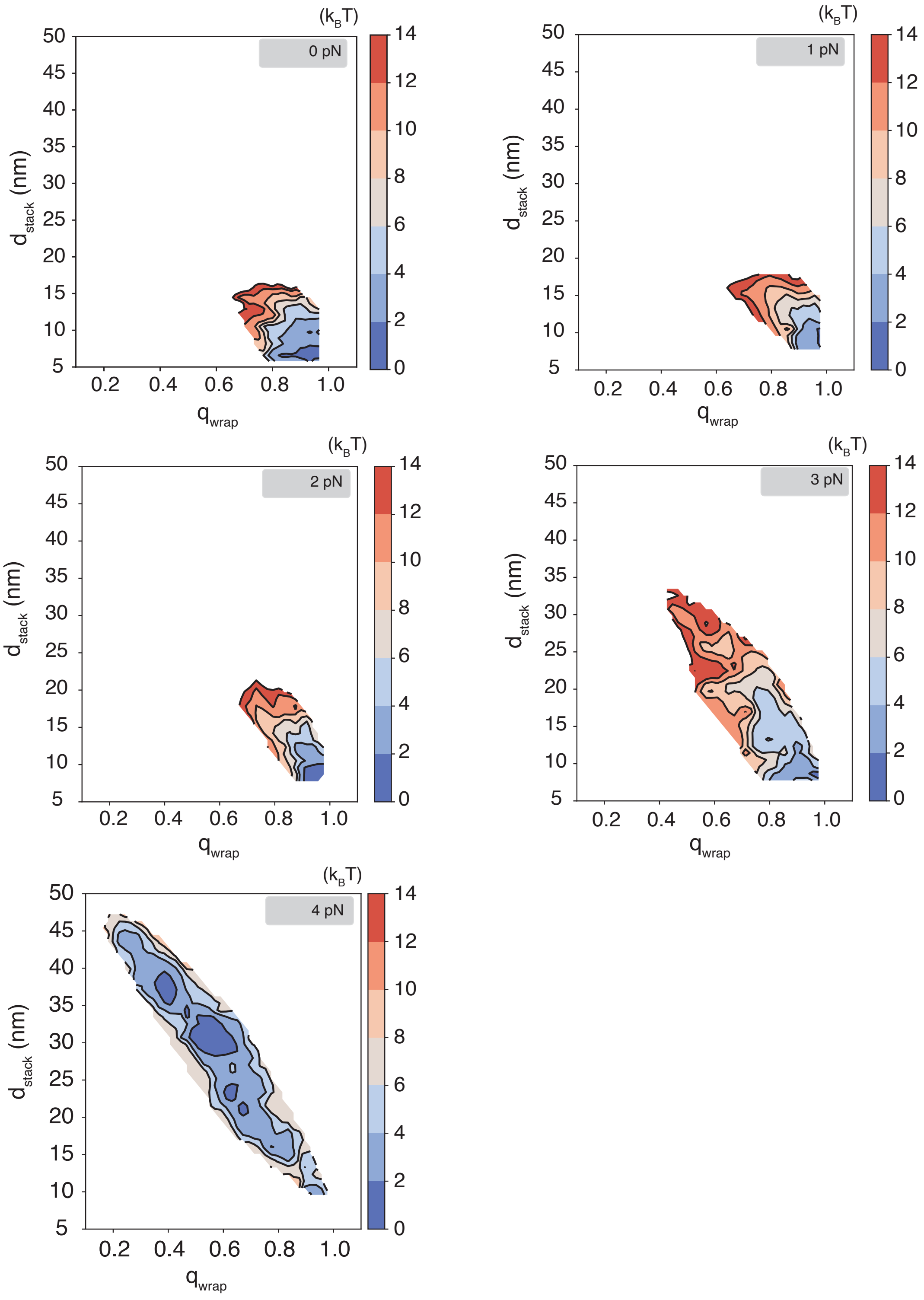


Figure S1: Two dimensional free energy profiles as a function of nucleosome unwrapping ($q_{\mathrm{wrap}}$) and unstacking ($d_{\mathrm{stack}}$) at various extension forces determined from umbrella simulations. See text *Section: Free Energy Profiles for Chromatin Under Tension* for simulation details.

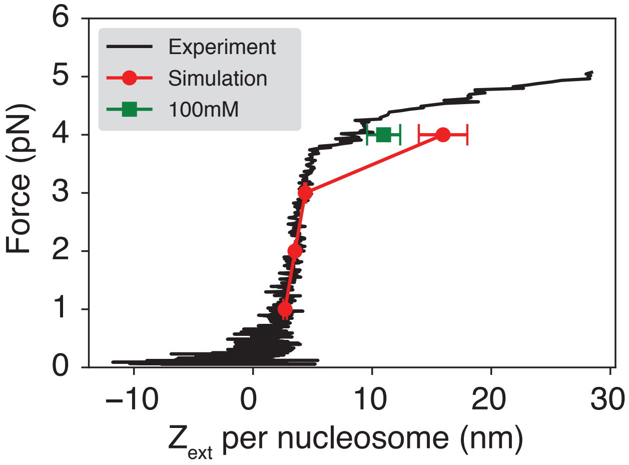


Figure S2: Comparison between the simulated (red) and experimental[S24] (black) force-extension curves. The results for simulations performed with 150 mM monovalent ions are reproduced from Fig. 1B. The green dot corresponds to chromatin extension at 4pN force obtained from simulations with 100 mM monovalent ions.

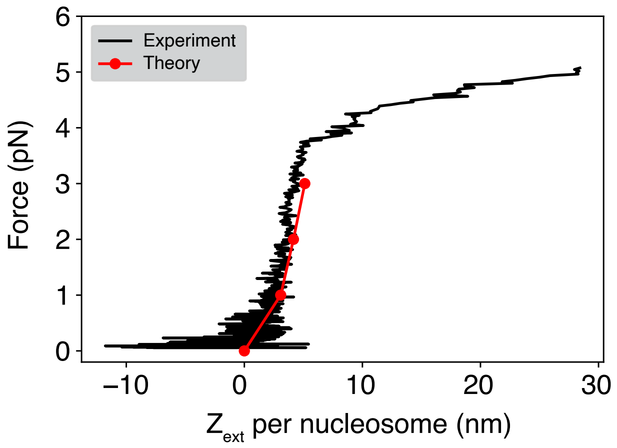


Figure S3: Theoretical predictions of chromatin extension along the $z$ axis, $Z_{\text{ext}}$. We assumed a harmonic potential for the end-to-end distance of the unbiased chromatin. Parameters in the potential were obtained from a least square fitting to the simulation results shown in Figure 1C at 0 pN. From the harmonic potential, $Z_{\text{ext}}$ can be computed with the analytical expression provided in Eq. S9. See *Section: Theoretical predictions of chromatin extension along the z axis* for a detailed discussion.

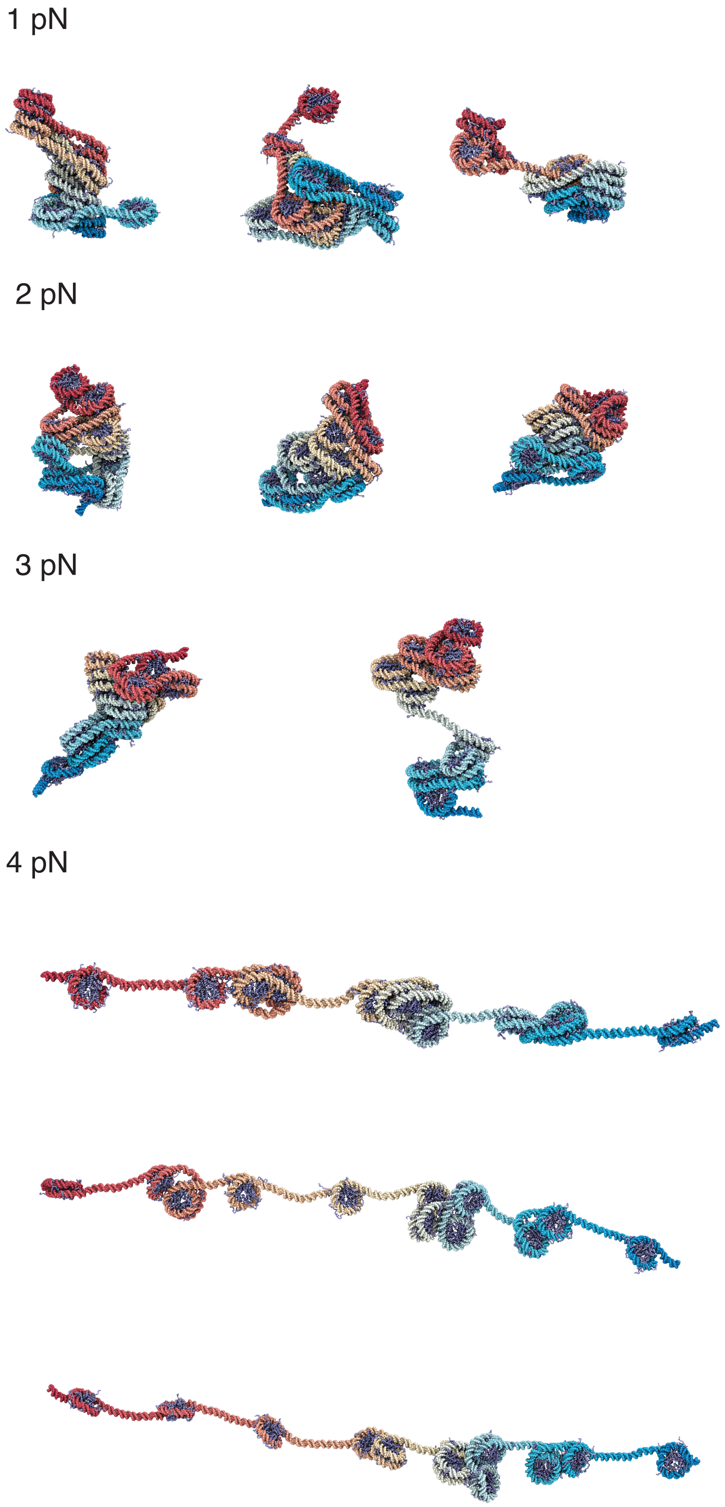


Figure S4: Additional representative chromatin structures from simulations performed under various extension forces. The values for the extension force are provided next to the structures. Similar to the ones shown in Figure 2 of the main text, these structures correspond to the central configurations of the clusters identified by the single-linkage algorithm using root mean squared distance (RMSD) as the distance between structures.

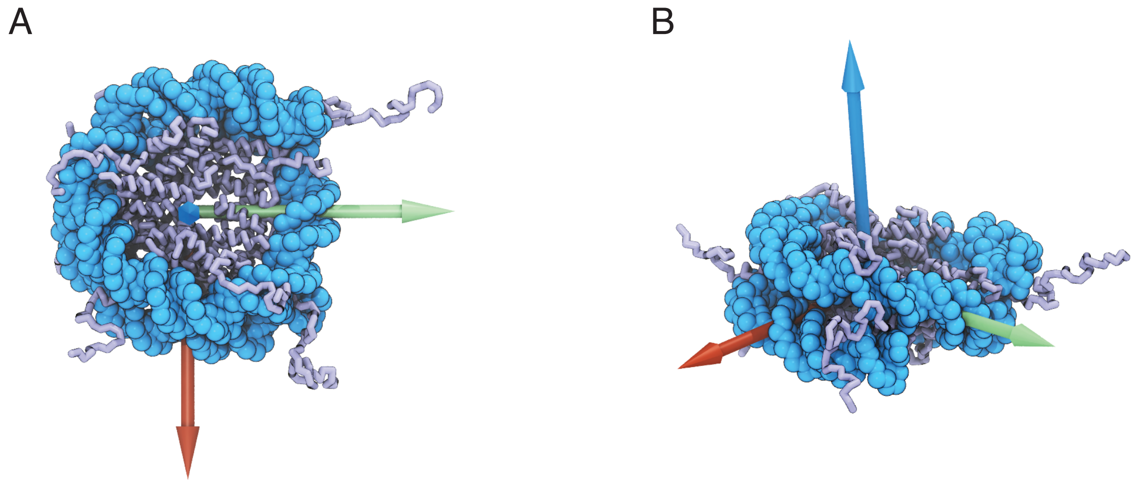


Figure S5: Illustration of the nucleosome coordinate system used to distinguish shearing and normal motions. The nucleosome is shown in the coarse-grained representation derived from the crystal structure (PDB ID: 1KX5).[S8] The origin of the coordinate system is defined as the center of residues 63-120, 165-217, 263-324, 398-462, 550-607, 652-704, 750-811, and 885-949. The red arrow points from the origin to the center of residues 63-120, 165-217, 750-811 and 885-949. The green arrow points towards the nucleosome dyad defined as the center of residues 81-131 and 568-618. The blue arrow is defined as the cross product of vectors along the red and the green arrows. See text *Section: Decomposing Inter-nucleosome distances into Shear and Normal Motion* for further discussions.

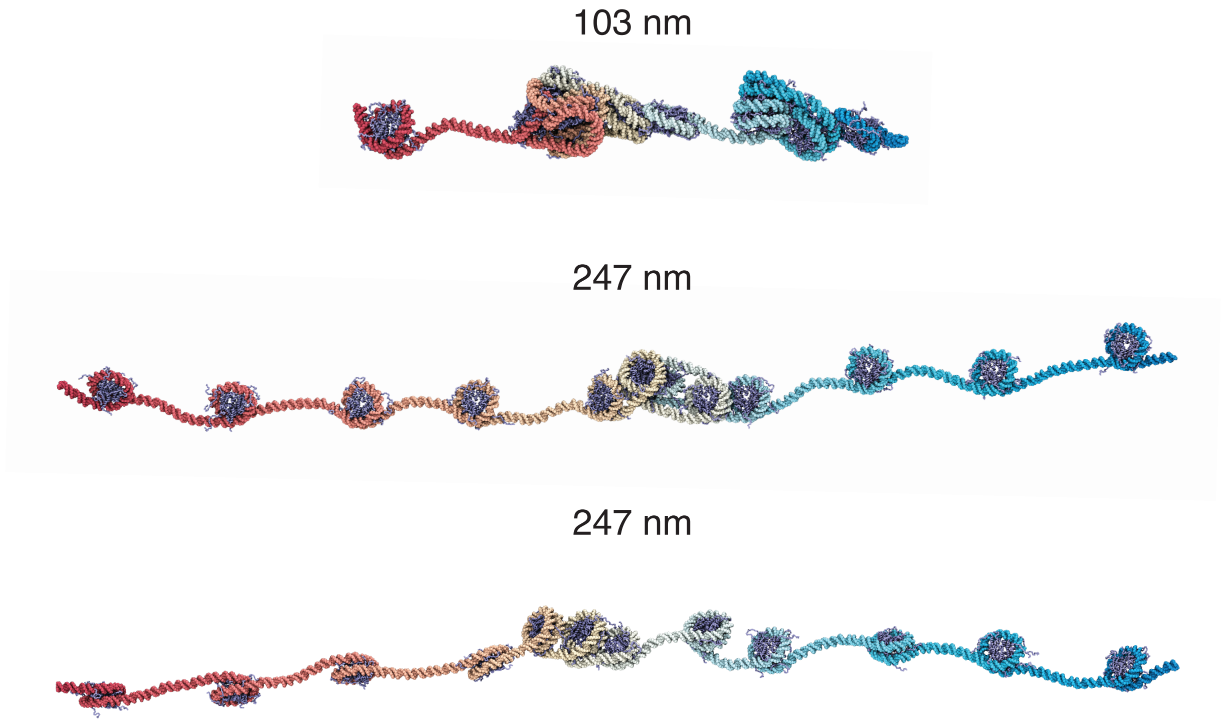


Figure S6: Additional representative chromatin structures at smaller and larger distances than the average extension at 4 pN force. The end-to-end distance are provided next to the structures.

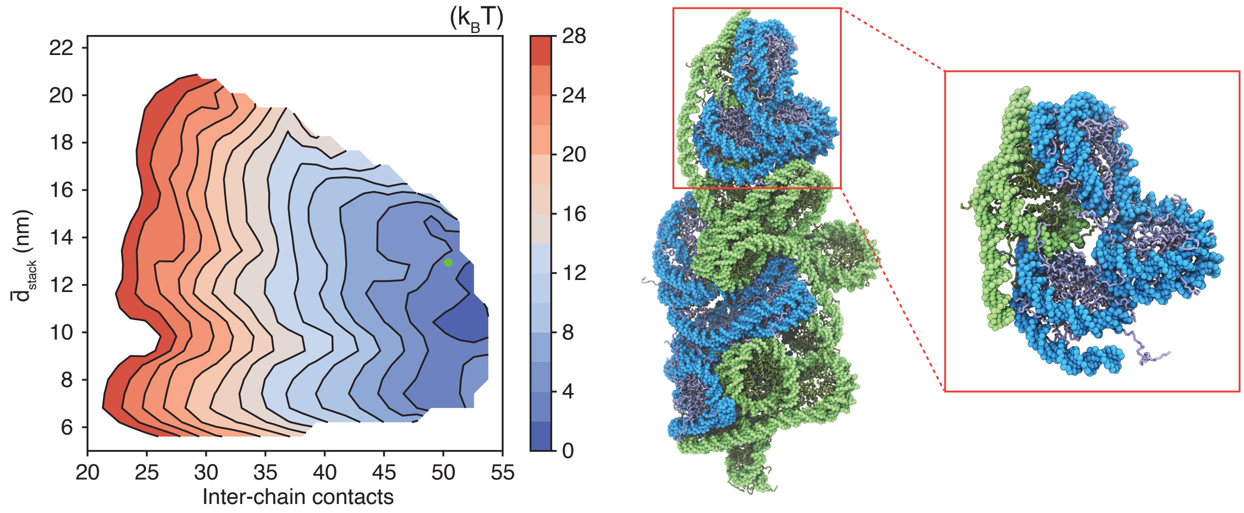


Figure S7: Representative structure of two contacting chromatin segments that adopt more extended configurations. Extension leads to more interdigitation between the two chains. The inset highlights the interactions between inter-chain nucleosomes. The free energy and collective variable values is indicated as the green dot in the free energy profile.

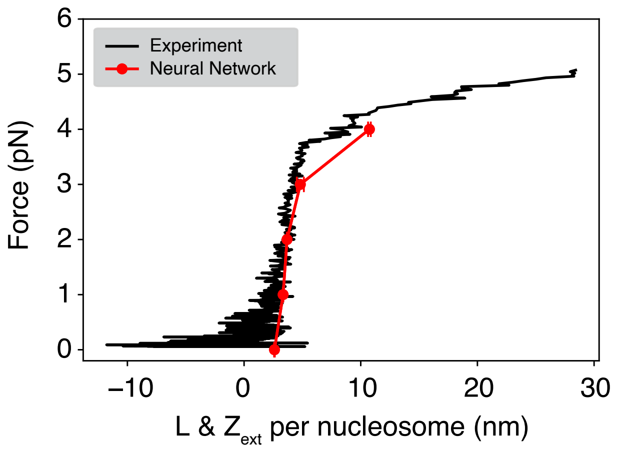


Figure S8: Comparison between experimental[S24] force-extension curve (black) and the one predicted by the neural network model. The neural network model quantifies chromatin stability as a function of inter-nucleosome distances. Based on the derivation shown in Eq. S9, when extension force is larger than 1 pN, the extension along $z$ axis ($L_z$) is very close to the end-to-end distance ($L$), so that we approximated the $z$-axis extension per nucleosome using the distance between first and last nucleosome ($L$) divided by 11. $L$ at different extension force was calculated using umbrella simulations of the neural network model. See text *Section: Initial configurations from the neural network model* and *Section: Neural Network Model for the 12mer Chromatin* for simulation details.

(S1) Plimpton, S. Fast Parallel Algorithms for Short-Range Molecular Dynamics. *Journal of Computational Physics* **1995**, *117*, 1–19.

(S2) Bonomi, M.; Branduardi, D.; Bussi, G.; Camilloni, C.; Provasi, D.; Raiteri, P.; Donadio, D.; Marinelli, F.; Pietrucci, F.; Broglia, R. A.; Parrinello, M. PLUMED: A Portable Plugin for Free-Energy Calculations with Molecular Dynamics. *Computer Physics Communications* **2009**, *180*, 1961–1972.

(S3) Kumar, S.; Rosenberg, J. M.; Bouzida, D.; Swendsen, R. H.; Kollman, P. A. The Weighted Histogram Analysis Method for Free-Energy Calculations on Biomolecules. I. The Method. *J. Comput. Chem.* **1992**, *13*, 1011–1021.

(S4) Noel, J. K.; Levi, M.; Raghunathan, M.; Lammert, H.; Hayes, R. L.; Onuchic, J. N.; Whitford, P. C. SMOG 2: A Versatile Software Package for Generating Structure-Based Models. *PLoS Comput Biol* **2016**, *12*, e1004794.

(S5) Ding, X.; Vilseck, J. Z.; Brooks III, C. L. Fast Solver for Large Scale Multistate Bennett Acceptance Ratio Equations. *J. Chem. Theory Comput.* **2019**, *15*, 799–802.

(S6) Song, F.; Chen, P.; Sun, D.; Wang, M.; Dong, L.; Liang, D.; Xu, R.-M.; Zhu, P.; Li, G. Cryo-EM Study of the Chromatin Fiber Reveals a Double Helix Twisted by Tetranucleosomal Units. *Science* **2014**, *344*, 376–380.

(S7) Schalch, T.; Duda, S.; Sargent, D. F.; Richmond, T. J. X-Ray Structure of a Tetranucleosome and Its Implications for the Chromatin Fibre. *Nature* **2005**, *436*, 138–141.

(S8) Davey, C. A.; Sargent, D. F.; Luger, K.; Maeder, A. W.; Richmond, T. J. Solvent Mediated Interactions in the Structure of the Nucleosome Core Particle at 1.9 Å Resolution. *J. Mol. Biol.* **2002**, *319*, 1097–1113.

(S9) Lu, X.-J. 3DNA: A Software Package for the Analysis, Rebuilding and Visualization of

Three-Dimensional Nucleic Acid Structures. *Nucleic Acids Research* **2003**, *31*, 5108–5121.

(S10) Michaud-Agrawal, N.; Denning, E. J.; Woolf, T. B.; Beckstein, O. MDAnalysis: A Toolkit for the Analysis of Molecular Dynamics Simulations. *J Comput Chem* **2011**, *32*, 2319–2327.

(S11) Gowers, R.; Linke, M.; Barnoud, J.; Reddy, T.; Melo, M.; Seyler, S.; Domański, J.; Dotson, D.; Buchoux, S.; Kenney, I.; Beckstein, O. MDAnalysis: A Python Package for the Rapid Analysis of Molecular Dynamics Simulations. Python in Science Conference. Austin, Texas, 2016; pp 98–105.

(S12) Theobald, D. L. Rapid Calculation of RMSDs Using a Quaternion-Based Characteristic Polynomial. *Acta Crystallogr A* **2005**, *61*, 478–480.

(S13) Liu, P.; Agrafiotis, D. K.; Theobald, D. L. Fast Determination of the Optimal Rotational Matrix for Macromolecular Superpositions. *J. Comput. Chem.* **2010**, *31*, 1561–1563.

(S14) Koslover, E. F.; Fuller, C. J.; Straight, A. F.; Spakowitz, A. J. Local Geometry and Elasticity in Compact Chromatin Structure. *Biophys J* **2010**, *99*, 3941–3950.

(S15) Wales, D. J. *Energy Landscapes*; Cambridge Molecular Science; Cambridge University Press: Cambridge, UK ; New York, 2003.

(S16) Ding, X.; Lin, X.; Zhang, B. Stability and Folding Pathways of Tetra-Nucleosome from Six-Dimensional Free Energy Surface. *Nat. Commun.* **2021**, *12*, 1–9.

(S17) Freeman, G. S.; Hinckley, D. M.; Lequieu, J. P.; Whitmer, J. K.; de Pablo, J. J. Coarse-Grained Modeling of DNA Curvature. *The Journal of Chemical Physics* **2014**, *141*, 165103.

(S18) Clementi, C.; Nymeyer, H.; Onuchic, J. N. Topological and Energetic Factors: What Determines the Structural Details of the Transition State Ensemble and "En-Route" Intermediates for Protein Folding? An Investigation for Small Globular Proteins. *J Mol Biol* **2000**, *298*, 937–953.

(S19) Noel, J. K.; Whitford, P. C.; Onuchic, J. N. The Shadow Map: A General Contact Definition for Capturing the Dynamics of Biomolecular Folding and Function. *J. Phys. Chem. B* **2012**, *116*, 8692–8702.

(S20) Miyazawa, S.; Jernigan, R. L. Estimation of Effective Interresidue Contact Energies from Protein Crystal Structures: Quasi-Chemical Approximation. *Macromolecules* **1985**, *18*, 534–552.

(S21) Zhang, B.; Zheng, W.; Papoian, G. A.; Wolynes, P. G. Exploring the Free Energy Landscape of Nucleosomes. *J. Am. Chem. Soc.* **2016**, *138*, 8126–8133.

(S22) Meng, H.; Andresen, K.; Van Noort, J. Quantitative Analysis of Single-Molecule Force Spectroscopy on Folded Chromatin Fibers. *Nucleic Acids Res.* **2015**, *43*, 3578–3590.

(S23) Moller, J.; Lequieu, J.; de Pablo, J. J. The Free Energy Landscape of Internucleosome Interactions and Its Relation to Chromatin Fiber Structure. *ACS Cent. Sci.* **2019**, *5*, 341–348.

(S24) Kaczmarczyk, A.; Meng, H.; Ordu, O.; van Noort, J.; Dekker, N. H. Chromatin Fibers Stabilize Nucleosomes under Torsional Stress. *Nat. Commun.* **2020**, *11*, 1–12.
